# Vertical HIV-1 transmission in the setting of maternal broad and potent antibody responses

**DOI:** 10.1101/2022.02.03.479075

**Authors:** Joshua J Tu, Amit Kumar, Elena E Giorgi, Josh Eudailey, Celia C LaBranche, David R Martinez, Genevieve G Fouda, Yvetane Moreau, Allison Thomas, David Montefiori, Feng Gao, Manish Sagar, Sallie R Permar

**Affiliations:** Duke Human Vaccine Institute, Duke University Medical Center, Durham, NC 27710, USA; BioAgilytix Labs, Durham, NC 27713, USA; Theoretical Division, Los Alamos National Laboratory, Los Alamos, NM 87544, USA; Department of Pediatrics, Weill Cornell Medicine, New York, NY 10065, USA; Department of Surgery, Duke University School of Medicine, Durham, NC 27710, USA; Department of Epidemiology, University of North Carolina at Chapel Hill, Chapel Hill, NC 27599, USA; Department of Molecular Genetics and Microbiology, Duke University Medical Center, Durham, NC 27710, USA; Department of Pediatrics, Duke University Medical Center, Durham, NC 27710, USA; Department of Medicine, Boston Medical Center, Boston University School of Medicine, Boston, Massachusetts, USA; Department of Medicine, Duke University Medical Center, Durham, NC 27710, USA; School of Medicine, Jinan University, Guangzhou, Guangdong 510632, P.R. China

## Abstract

Despite the worldwide availability of antiretroviral therapy (ART), approximately 150,000 pediatric HIV infections continue to occur annually. ART can dramatically reduce HIV mother-to-child transmission (MTCT), but inconsistent drug access and adherence, as well as primary maternal HIV infection during pregnancy and lactation are major barriers to eliminating vertical HIV transmission. Thus, immunologic strategies to prevent MTCT, such as an HIV vaccine, will be required to attain an HIV-free generation. A primary goal of HIV vaccine research has been to elicit broadly neutralizing antibodies (bnAbs) given the ability of passive bnAb immunization to protect against sensitive strains, yet we previously observed that HIV-transmitting mothers have more plasma neutralization breadth than non-transmitting mothers. Additionally, we have identified infant transmitted/founder (T/F) viruses that escape maternal bnAb responses. In this study, we examine a cohort of postpartum HIV-transmitting women with neutralization breadth to determine if certain maternal bnAb specificities drive the selection of infant T/F viruses. Using HIV pseudoviruses that are resistant to neutralizing antibodies targeting common bnAb epitopes, we mapped the plasma bnAb specificities of this cohort. Significantly more transmitting women with plasma bnAb activity had a mappable plasma bnAb specificity (six of seven, or 85.7%), compared to that of non-transmitting women with plasma bnAb activity (seven of twenty-one, or 33.3%, p=0.029 by 2-sided Fisher exact test). Our study suggests that having multispecific broad activity and/or uncommon epitope-specific bnAbs in plasma may be associated with protection against the vertical HIV transmission in the setting of maternal bnAb responses.

**Importance:** As mother to child transmission (MTCT) of HIV plays a major part in the persistence of the HIV/AIDS epidemic and bnAb-based passive and active vaccines are a primary strategy for HIV prevention, research in this field is of great importance. While previous MTCT research has investigated the neutralizing antibody activity of HIV-infected women, this is to our knowledge the largest study identifying differences in bnAb specificity of maternal plasma between transmitting and non-transmitting women. Here, we show that among HIV-infected women with broad and potent neutralization activity, more postpartum-transmitting women had a mappable plasma broadly neutralizing antibody (bnAb) specificity, compared to that of non-transmitting women, suggesting that the non-transmitting women more often have multi-specific bnAb responses or bnAb responses that target uncommon epitopes. Such responses may be required for protection against vertical HIV transmission in the setting of maternal bnAb responses.

## Introduction

Antiretroviral therapy (ART) can dramatically reduce mother-to-child transmission (MTCT) of HIV-1, but challenges remain in ART access and adherence during pregnancy and throughout breastfeeding. Despite the worldwide availability of ART, around 150,000 new pediatric HIV infections occurred in 2020 (1). Further, with interrupted ART supply and access during the Severe Acute Respiratory Syndrome-Coronavirus-2 (SARS-CoV-2) pandemic that began in 2019, this number is likely to have increased significantly (2). Moreover, acute maternal infections during pregnancy and breastfeeding are not addressed in the current ART-based strategies to reduce or eliminate MTCT. Thus, additional strategies to prevent MTCT will be required to attain an HIV-free generation. Interestingly, in the absence of ART, only 15-30% of HIV-infected mothers transmit the virus to their child with additional risk associated with breastfeeding, suggesting that there are immunologic factors that modify the risk of MTCT of HIV-1 (3, 4). Maternal vaccination strategies seek to enhance immune factors that are partially protective against vertical HIV transmission to further reduce HIV MTCT.

To design immune-based strategies that can synergize with ART to further reduce MTCT, it is important to understand the role of circulating and placentally transferred maternal antibodies in vertical transmission. Many studies have shown that despite having a diverse array of HIV-1 variants circulating in the mother (5–10), only one or a few maternal variants get transmitted to the infant, known as the infant transmitted/founder (T/F) variant(s). This genetic bottleneck suggests that the maternal immune system selects for viruses that are transmitted to the infants. Yet, while broadly neutralizing antibodies (bnAbs) have been shown to protect against susceptible virus strains in animal studies (11–16), there is no consensus about the role of neutralizing antibodies (nAbs) in driving the genetic bottleneck of MTCT (17–19). We previously studied the role of maternal antibodies in the selection of infant T/F viruses with a cohort of 16 clade B infected, peripartum transmitting mother-infant pairs from the United States (17) and demonstrated that infant T/F viruses are more neutralization-resistant to paired maternal plasma compared to non-transmitted maternal variants. Our findings were in line with a previous study investigating autologous virus neutralization activity in 12 transmission pairs from Nairobi, Kenya (19), yet resistance to maternal plasma neutralization was not a defining feature of infant T/Fs in another study of 10 *in utero* and 9 intrapartum, clade C-infected pairs from Malawi (18). Differences in the results of these studies could be due to sample size, HIV-1 subtype, and mode of transmission. Hence, it remains unclear if maternal plasma neutralizing activity drives infant T/F selection.

Conflicting findings have also been observed in studies comparing the plasma neutralization activity of transmitting and non-transmitting mothers against circulating, non-transmitted maternal variants. Some studies have shown that autologous virus neutralizing activity is higher in non-transmitting women than transmitting women (8, 20), while more recent studies observed no difference at all (18, 21). In addition to autologous virus neutralization, some studies have tested heterologous virus neutralization activity of maternal and infant plasma as a risk factor for MTCT. For example, in a study of postpartum transmission, breadth and potency of infant plasma were tested against a global panel of eight heterologous viruses (22). No correlation of heterologous virus breadth and potency of placentally acquired neutralizing antibodies with transmission was observed. In contrast, we previously reported that HIV-transmitting mothers had significantly higher heterologous virus neutralization breadth and potency than that of non-transmitting mothers against a different global panel of 11 viruses in a cohort of 63 mother-infant pairs from the Breastfeeding, Antiretrovirals, and Nutrition (BAN) study of breast milk transmission (23, 24). High breadth and potency of maternal plasma suggests that the development of maternal bnAb activity in plasma may be associated with risk of MTCT, a mechanism of which is yet unknown.

BnAb activity in plasma has been identified in about 10-25% of chronically HIV-infected adults and targets highly conserved epitopes on the HIV envelope, allowing them to neutralize a majority of HIV variants found globally with high potency (25–30). Additionally, most chronically infected individuals display low-to-moderate plasma neutralization breadth (31). Since MTCT occurs in the presence of maternal antibodies, it is important to understand the role of maternal bnAbs in the prevention of MTCT, as well as the impact of future bnAb-based prophylaxis and therapy approaches on MTCT. Viral evolution and escape from naturally occurring bnAbs in HIV-infected individuals has been well characterized (32–36), yet while bnAb-resistant infant T/Fs have been identified (37–40), the selection and escape of such viruses has not previously been studied in relationship to maternal bnAb activity. Importantly, we recently identified an infant T/F virus that was distinguished from non-transmitted maternal variants by resistance to V3 glycan-specific plasma bnAb responses in paired maternal plasma (41), indicating that maternal bnAb activity can drive the selection of escape variants leading to MTCT.

Taken together, broad and potent neutralization activity of maternal plasma is clearly not sufficient for protection against MTCT. Yet, with passive immunization and induction of such plasma neutralization activity as a current dominant approach to preventing HIV acquisition, it is imperative to identify whether there are specific characteristics of broad and potent neutralization responses that select for vertically transmitted variants. In this study, we characterized and mapped the bnAb specificity of maternal plasma from transmitting and non-transmitting mothers using samples selected from two HIV-infected maternal-infant cohorts prior to the era of universal ART in pregnancy (Breastfeeding, Antiretroviral, and Nutrition Study and the Center for HIV/AIDS Vaccine Immunology 009, both in Malawi) using the criteria of broad and potent neutralization activity of plasma against a global virus panel (23). We found that transmitting mothers more often have mappable single epitope-specific bnAb activity in plasma than non-transmitting mothers, suggesting that polyclonal or uncommon neutralization responses occur more frequently in non-transmitting women. Overall, these findings will help define the characteristics of protective neutralizing antibody responses against MTCT, bringing insight to bnAb-based HIV vaccine development and the elimination of pediatric HIV.

## Methods

### Study Design

Patient samples were selected from the Breastfeeding, Antiretroviral, and Nutrition Study (BAN) and the Center for HIV/AIDS Vaccine Immunology 009 (CHAVI009) cohorts, both enrolled in Malawi in 2004-2010 and 2007-2009, respectively. Both cohorts were infected primarily with clade C HIV, and their enrollment criteria have been previously described (24, 37, 42, 43)(ClinicalTrials.gov no. NCT00164736). BAN cohort samples were selected from the control arm and received a combination ART for a duration of 7 days after delivery (44). CHAVI009 cohort patients received one dose of the antiviral Nevirapine at the time of delivery. Transmission events were identified by HIV DNA PCR testing of the infants at birth, at 4-6 weeks, and every 3 months until weaning. The CHAVI009 study was approved by the College of Medicine Research and Ethics Committee in Malawi and by all participating institutions’ institutional review boards (IRBs). The BAN Study was approved by the Malawi National Health Science Research Committee and by all participating institutions’ IRBs. All women provided written informed consent for themselves as well as on behalf of their infants. Maternal and infant samples from the CHAVI009 and BAN cohorts were received as deidentified material and were deemed “not human subject research” by the Duke University School of Medicine IRB (BAN: Duke IRB Pro00030437, CHAVI009: Duke IRB Pro00016627) and Boston University School of Medicine IRB.

### Neutralization Assays

Maternal plasma and monoclonal antibodies were tested against the 11 virus global panel and infant T/Fs pseudoviruses for neutralization activity using a TZM-bl neutralization assay as described previously (23). Inhibitory dilution 50% (ID_50_)s) and inhibitory concentration 50% (IC50s) were measured for maternal plasma or monoclonal antibodies using 3-fold dilutions ranging from 1:40 to 1:87480 and 25 μg/mL to 0.011 μg/mL respectively. To identify HIV-1 Env epitopes targeted by the neutralizing antibodies present in the maternal plasma, we assessed neutralization against HIV Env pseudoviruses with unique mutations in established bnAb epitopes, including: CD4 binding site (TRO.11.N279A, TRO.11.G458Y, TRO.11.N276Q, X1632.D279N, X1632.G458Y, and X1632.N276Q), V2 glycan (TRO.11.N160K and BJOX2000.N160K), V3 glycan (TRO.11.N332A and BJOX2000.N332A), and MPER (TRO.11.W672A and BJOX2000.W672A). Neutralization epitope-specificity was identified as a 2 fold reduction or greater in ID_50_ of a mutant pseudovirus compared to wild type.

### Screening Maternal Plasma for Broad and Potent Neutralizing Activity

Maternal plasma samples collected both prior and closest to the first positive infant HIV test (27-66 days) from the BAN cohort were previously tested for breadth, and the plasma that neutralized TRO11 and BJOX2000 (2 viruses from the global panel) at an ID_50_ ≥120 were selected from the available samples (n = 6) (24). To match the transmitting mother sample timepoints, 67 plasma samples collected 4-6 weeks after delivery from mothers from the CHAVI009 cohort who only received peripartum ART were screened for neutralization against 11 global panel HIV Env pseudovirus (23, 37, 42, 43). Broad neutralization activity was defined as plasma neutralization at an ID_50_ ≥50 for six of the eleven viruses tested. For samples not tested at a dilution of 50 in Figure 2A, we calculated the % neutralization at a dilution of 50 using the mathematical formulation for a single antibody neutralization curve(45). The breadth and potency (BP) score was defined according to equation 1 in Ghulam-Smith et al. (24).

### Pseudovirus Preparation

HIV-1 Env pseudoviruses were produced as previously described (46) in 293T cells through co-transfection using 4 μg of envelope expression plasmid and 8 μg of backbone plasmid pSG3ΔEnv or pQ23ΔEnv. VSV-G and SVA.MulV viruses were prepared by co-transfection in 293T cells with backbone plasmid pNL-LucR-E and pSG3ΔEnv respectively. Virus was harvested after 48 hours post transfection, filtered and stored at −80°C at 20% FBS concentration. Pseudoviruses were tittered by infecting Tzm-bl cells at 5-fold dilutions from 1:10 to 1:97,656,250 and the dilution that achieves 50,000 relative luminescence units (RLU) during infection was used in all assays. If the pseudovirus infection did not reach 50,000 RLU, then a dilution that provided an RLU ten times higher than the uninfected cell control was chosen for assays. Maternal amplicons for pseudovirus preparation were selected using a previously used algorithm to represent viral diversity (37). 2315 infant amplicons for pseudovirus preparation were visually selected based on the unique mutations in the known bnAb epitopes (Fig. 5C).

### Monoclonal antibody production

Monoclonal antibodies PGT121, PG9, VRC01, and 10E8 were produced with plasmids obtained through the NIH HIV Reagent Program, Division of AIDS, NIAID, NIH. Heavy and light chain plasmids were transfected into 293i cells using the Expifectamine kit (ThermoFisher A14524) as per manufacturer guidelines. Transfected cells were incubated for 5 days before harvesting the cell supernatant. Antibodies were purified from culture supernatants using affinity chromatography with protein A beads as previously described (47). After purification, all antibodies were then buffer exchanged into a citric acid buffer (pH 6) before use.

### VRC01-blocking ELISA

VRC01-blocking activity of maternal plasma was detected by competition ELISA method as previously described (41). Briefly, 384 well plates were coated with 45 ng/mL of 1086CK160N gp120 overnight at 4°C. 10ul of plasma diluted at 1:30 were added in duplicate and incubated at RT for one hour. Plates were washed twice and 16ng of biotinylated VRC01 was added and incubated at RT for one hour. Plates were washed twice, and HRP-conjugated streptavidin was used at 1:10000 dilution and incubated at RT for one hour. Plates were washed four times and developed using 20μl of SureBlue substrate per well and incubated for 3 minutes and 30 secs. The reaction was stopped using stop solution and the plate was read using a spectrophotometer at 450 nm. Percent blocking was determined by the percent difference in OD (max OD=2.23) between uninhibited bnAb binding and the sample. 50% blocking was defined as the positivity cutoff.

### Single Genome Amplification (SGA)

SGA sequences were obtained as described previously (37). In brief, RNA was extracted from breast milk and plasma samples and reverse transcribed. The obtained cDNA was used to PCR amplify full length *env* genes. PCR reactions were performed in 96 well plates at a dilution of cDNA to yield no more than 20% PCR positive wells to fit a Poisson distribution of one template per reaction. Autologous *env* sequences were aligned using Seaview (48) and compared using LANL Gene Cutter (https://www.hiv.lanl.gov/content/sequence/GENE_CUTTER/cutter.html) and SeqPublish (https://www.hiv.lanl.gov/content/sequence/SeqPublish/seqpublish.html). Key broadly neutralizing antibody binding sites were found using LANL Neutralizing Antibody Contacts and Features database (https://www.hiv.lanl.gov/components/sequence/HIV/featuredb/search/env_ab_search_pub.comp) and identified in the autologous virus envelope sequences using LANL HIV Sequence Locator (https://www.hiv.lanl.gov/content/sequence/LOCATE/locate.html) to compare them to HXB2. Total envelope glycosylation count was calculated using LANL N-Glycosite (https://www.hiv.lanl.gov/content/sequence/GLYCOSITE/glycosite.html) and variable region length and glycosylation were calculated using LANL Variable Region Characteristics (https://www.hiv.lanl.gov/content/sequence/VAR_REG_CHAR/index.html).

### Statistical Analysis

Counts of mapped versus non-mapped bNAb specificities were compared between transmitting and non-transmitting mothers using a 2-sided Fisher exact test. Breadth and breadth-potency scores between transmitting and non-transmitting mothers were compared using 2-sided Wilcoxon tests. All tests were run on the R platform (https://www.R-project.org). Heatmaps were created using the Heatmap tool provided on the LANL database (https://www.hiv.lanl.gov/content/sequence/HEATMAP/heatmap_mainpage.html). Phylogenetic trees were created with MEGA-X using a Kimura 2 Parameters (K2P) model (49).

Env variable loop lengths, number of glycosylation sites, and net charge were obtained using the LANL tool Variable Region Characteristics (https://www.hiv.lanl.gov/content/sequence/VAR_REG_CHAR/index.html) and compared between mother and infant for ID 2315 using 2-sided Wilcoxon test.

## Results

### Identifying HIV-infected transmitting and non-transmitting mothers with plasma HIV-neutralization breadth

All transmission events occurred postnatally, with the infants testing HIV DNA PCR positive after 4-6 weeks of age. BAN cohort maternal plasma samples were previously assessed for breadth (24). Of 19 transmitting mothers from this cohort, 16 neutralized 6 or more viruses from the 11 virus global panel. Of the 44 non-transmitting mothers in the BAN cohort, 24 mothers neutralized 6 or more of the 11 viruses from the global panel. From the CHAVI009 cohort, we initially screened 3 transmitting and 62 non-transmitting mothers for neutralization breadth. Of the transmitting mothers, only one (subject 1209) demonstrated neutralization breadth. Of the 62 non-transmitting mothers in the CHAVI cohort, 18 demonstrated neutralization breadth. Combining the cohorts, we screened a total of 22 transmitting and 106 non-transmitting mothers, for a total size of 128 samples (Fig. 1). Consistent with previous findings that BAN cohort transmitting mothers had higher neutralizing breadth than non-transmitting mothers (p<0.001)(24), the combined cohort had more transmitting mothers (17/22, or 77%) than non-transmitting mothers (42/106, or 40%) that displayed neutralization breadth.

**Figure 1.**
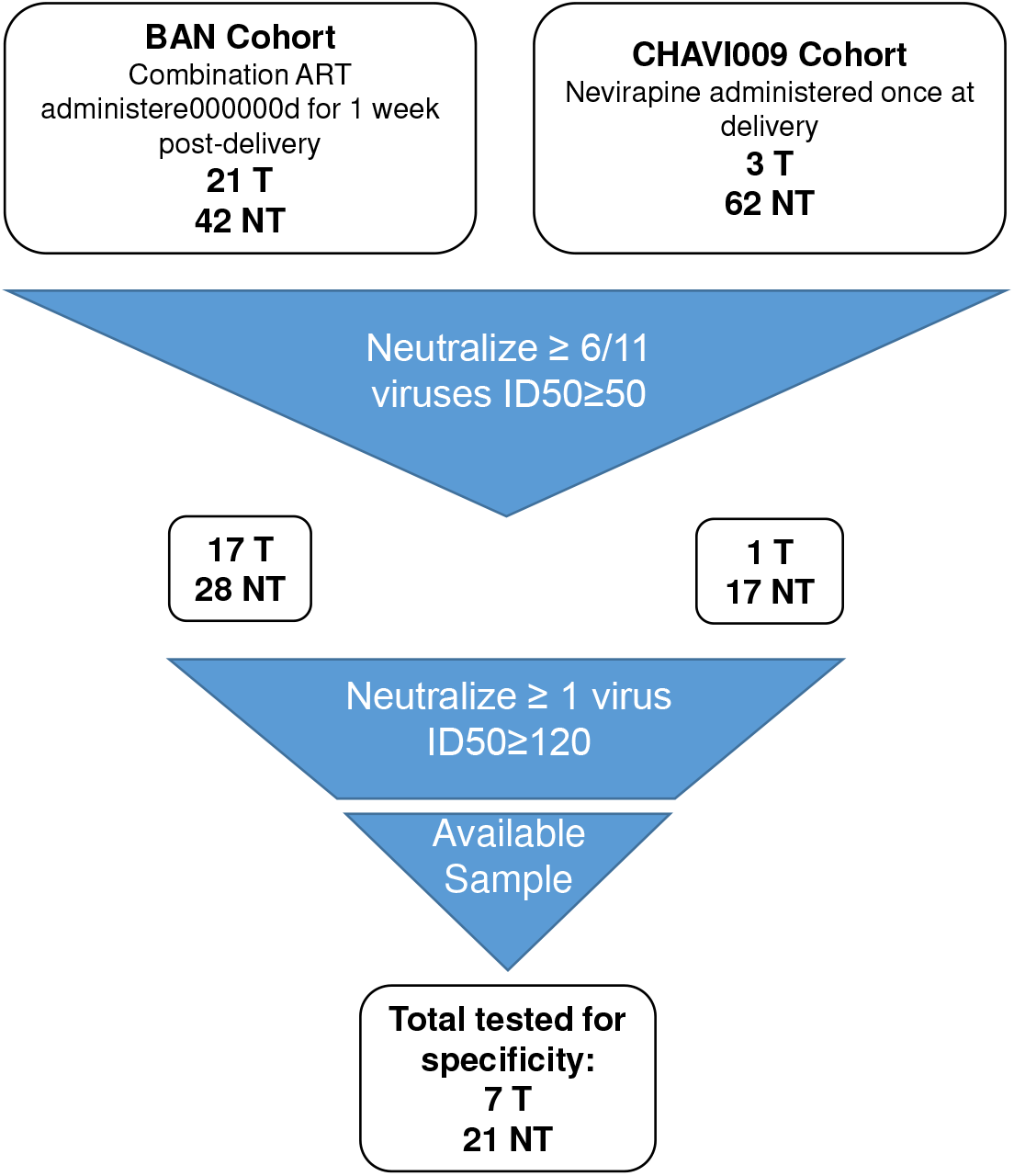
Diagram of the screening criteria used to identify mothers with bnAb activity. Transmitting and non-transmitting mothers were screened for plasma neutralization breadth and potency to determine if they had plasma broad neutralizing activity. Mothers with at least 140μl of available plasma and ID_50_ >120 against at least 1 virus met criteria for this study.

In order to explore the specificity of the plasma neutralizing activity in HIV-transmitting and nontransmitting women, we selected a subcohort of mothers with comparable breadth and neutralization potency. To this end, we selected plasma samples that had sufficient volume (>140 μl) and neutralized at least one virus with an ID_50_ >120 to facilitate neutralization epitope-specificity mapping (Fig. 1). This ID_50_ cut off was chosen based on the ability to observe a three-fold reduction in neutralization activity against a bnAb epitope mutant virus measured at a starting plasma dilution of 1:40. Six transmitting and four non-transmitting women from the BAN cohort and one transmitting and 17 non-transmitting women from the CHAVI009 cohort met criteria for the neutralization epitope mapping, yielding a study sample size of seven transmitting and 21 non-transmitting women (Fig 2A, Table 1). By design, given how we selected the women, the Breadth Potency Score (BP score, as defined by Ghulam-Smith et al.(24) and the number of viruses neutralized was not different (p=0.12 by Wilcoxon test) among the transmitting and non-transmitting women selected for inclusion in our analysis based on the selection criteria (Fig. 2B).

**Figure 2.**
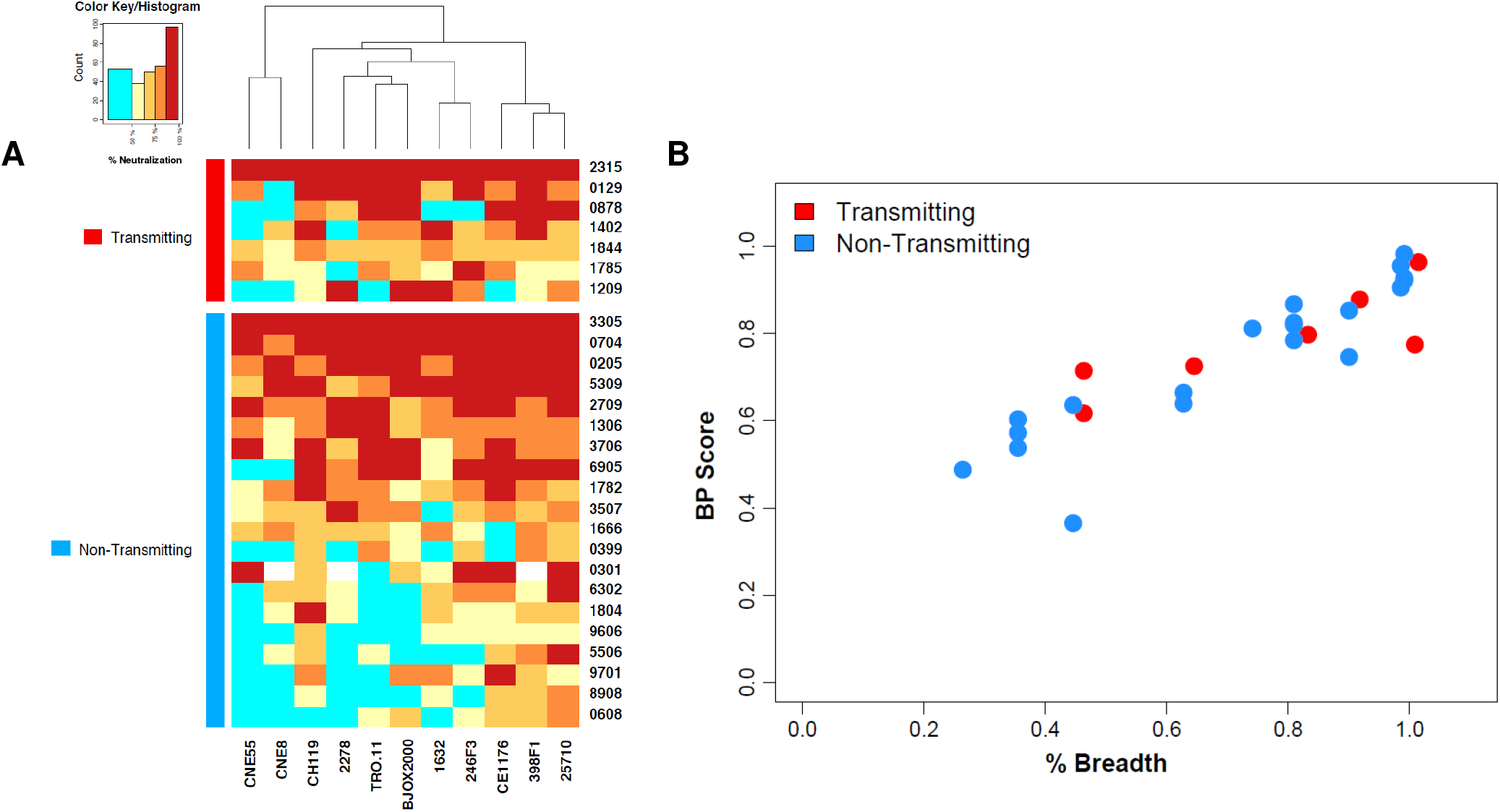
Breadth-potency comparison of transmitting and non-transmitting HIV-infected women with plasma neutralization breadth. A) Maternal plasma was tested against an 11-virus global panel. The % neutralization at a dilution of 1:50 is displayed in the heat map. Samples not tested at an 1:50 dilution have their % neutralization calculated for said dilution using the mathematical formulation for a single antibody neutralization curve. Colors for % neutralization are described on the color key/histogram. B) Plot of the % of viruses neutralized (x-axis) vs. breadth/potency score (y-axis) for transmitting (red) and non-transmitting (blue) women. The breadth/potency score was not significantly different in between transmitting and non-transmitting women in this cohort.

**Table 1:** Maternal clinical characteristics, timepoints of infant HIV diagnosis, and timepoints of maternal plasma samples of Malawian HIV-infected transmitting and non-transmitting women.

| Maternal ID | Days PP Sample Collection | Maternal CD4<sup>+</sup> cells/mm<sup>3</sup> | Maternal log<sub>10</sub> plasma VL | Days PP Infant 1st HIV<sup>+</sup> | # Viruses Neutralized Over 50% | Maternal ID | Days PP Sample Collection | Maternal CD4<sup>+</sup> cells/mm<sup>3</sup> | Maternal log<sub>10</sub> plasma VL | # Viruses Neutralized Over 50% |
| --- | --- | --- | --- | --- | --- | --- | --- | --- | --- | --- |
| 129 | 174 | 309 | 4.39 | 240 | 10 | 1306 | 14 | 360 | 4.43 | 11 |
| 878 | 13 | 248 | 5.02 | 56 | 7 | 1666 | 13 | 258 | 4.91 | 10 |
| 1402 | 14 | 293 | 5.64 | 56 | 9 | 1782 | 171 | 293 | 4.68 | 11 |
| 1785 | 168 | 336 | 4.29 | 196 | 10 | 399 | 84 | 396 | 4.39 | 6 |
| 1844 | 15 | 426 | 4.59 | 42 | 11 | 704 | 86 | 293 | 3.99 | 11 |
| 2315 | 1 | 312 | 4.78 | 43 | 11 | 205 | 32 | 156 | 2.6 | 11 |
| 1209 | 25 | 94 | 5.24 | 200 | 7 | 3305 | 118 | 284 | 5.43 | 11 |
|  |  |  |  |  |  | 6905 | 28 | 123 | 4.45 | 11 |
|  |  |  |  |  |  | 5309 | 33 | 434 | NA | 11 |
|  |  |  |  |  |  | 3507 | 84 | 385 | 3.9 | 9 |
|  |  |  |  |  |  | 3706 | 87 | 571 | 3.73 | 11 |
|  |  |  |  |  |  | 2709 | 29 | 166 | 2.6 | 11 |
|  |  |  |  |  |  | 9701 | 31 | 153 | 4.8 | 8 |
|  |  |  |  |  |  | 301 | 28 | 406 | 4.88 | 11 |
|  |  |  |  |  |  | 4707 | 32 | 247 | 4.9 | 7 |
|  |  |  |  |  |  | 1804 | 48 | 833 | 2.68 | 8 |
|  |  |  |  |  |  | 9606 | 31 | 558 | 3.71 | 6 |
|  |  |  |  |  |  | 6302 | 33 | 115 | NA | 8 |
|  |  |  |  |  |  | 5506 | 25 | 420 | NA | 6 |
|  |  |  |  |  |  | 8908 | 31 | 178 | 3.97 | 6 |
|  |  |  |  |  |  | 0608 | 60 | 275 | 2.6 | 6 |

Red: Transmitting
Black: Nontransmitting
PP: Postpartum
NA: Not Available

### Neutralization epitope specificity mapping

To identify potential differences in the predominant neutralization epitope specificity in transmitting and non-transmitting women with broad and potent neutralizing plasma responses, we further investigated the specificities of broad neutralization activity in both HIV-transmitting and non-transmitting mothers. Using HIV pseudoviruses that have mutations in epitopes targeted by bnAbs, we sought to map the bnAb specificity of maternal plasma with broad neutralization activity (Fig 3). The panel included viruses with mutations to the CD4 binding site (N279A or D279N, G458Y, and N276Q), the V2 glycan (N160K), the V3 glycan (N332A), and the membrane-proximal external region (W672A) (see Methods section). We were able to map 6 of 7 (85.7%) transmitting women’s plasma broad neutralizing activity, two to the N276 CD4 binding site (CD4bs), one to the N160 V2 glycan, and two to the N332 V3 glycan. One transmitting mother’s broad neutralizing plasma activity was mapped to both the N276 CD4bs and the V3 glycan residue. Of the non-transmitting mothers’ plasma, we were able to map 7 of 21 (33.3%) that had neutralizing activity. The plasma broad neutralizing activity of four of the non-transmitting mothers was mapped to the N276 CD4bs, while V2 glycan, and V3 glycan regions were mapped for one non-transmitting maternal plasma each. One non-transmitting mother’s plasma broad neutralizing activity was also mapped to both the N276 CD4bs and V3 glycan epitopes (Fig 3C). Yet, for the majority of non-transmitting mothers (n = 14, 66.6%), we were not able to map the plasma broad neutralizing activity to a bnAb epitope, indicating that their plasma neutralizing responses are either multispecific, or alternatively, target an epitope not included in the mutant virus panel. Transmitting mothers in our cohort more frequently had epitope-mappable plasma bnAb specificity that targets at least one of the 4 most common bnAb epitopes (OR=10.9, p=0.029, 2-sided Fisher exact test) compared to non-transmitting women (Fig. 3C). These results suggest that the plasma HIV-neutralizing breadth among transmitting versus non-transmitting mothers with broad and potent neutralizing activity may be mediated via different antibody specificities.

**Figure 3.**
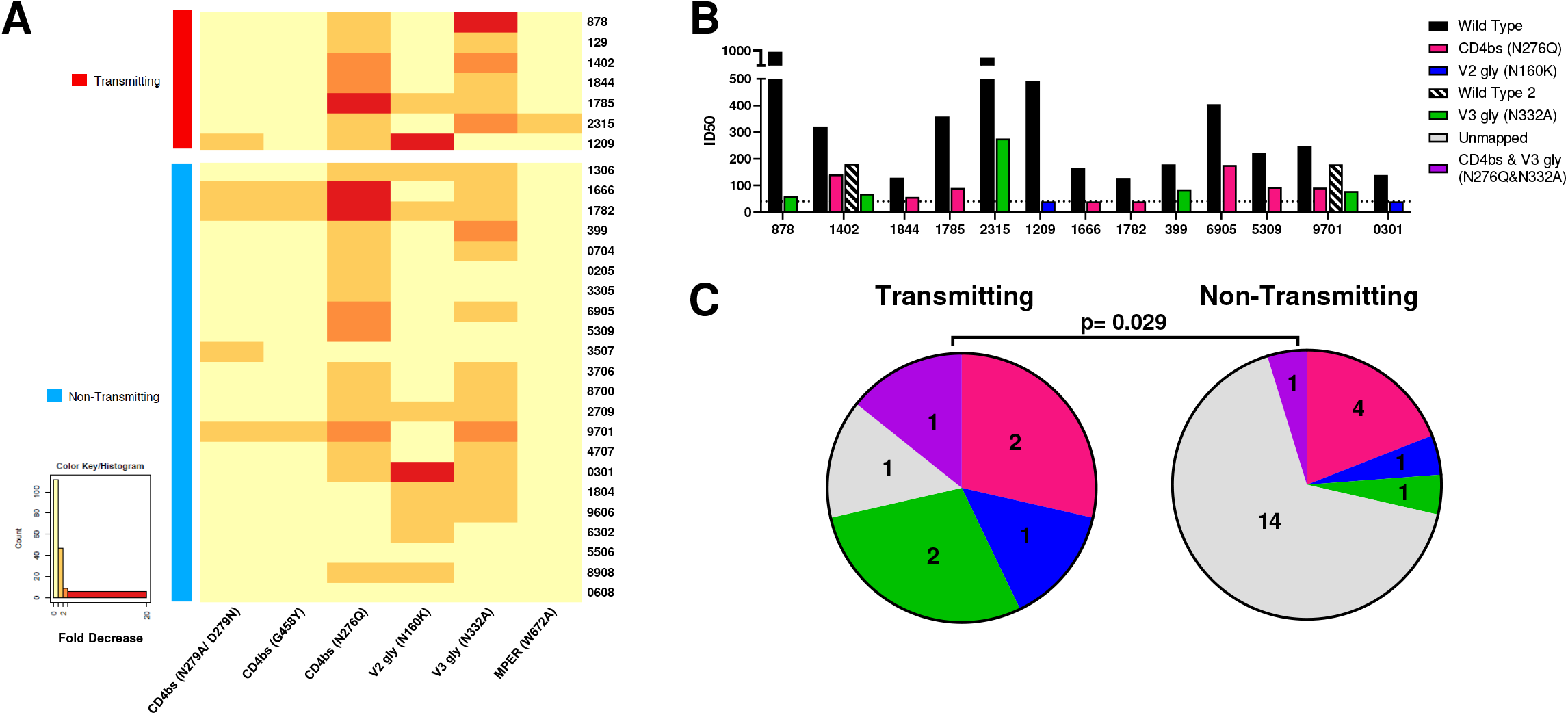
Plasma bnAb epitope-specificity was more often identified in HIV-infected transmitting women compared to non-transmitting women. A) Mothers with plasma broad neutralizing activity were tested for bnAb specificities using mutant envelope (Env) pseudoviruses (pseudovirus). Activity was measured by fold change compared to the wild type Env pseudovirus. Colors for fold decrease are described on the color key/histogram. A >2 fold decrease in neutralization compared to the wild type pseudovirus was our positivity threshold. B) ID_50_s for wild type and mutant pseudoviruses are shown for women with mapped specificities. C) Mapping data is summarized as a pie chart. Significantly more transmitting women were able to be mapped to a bnAb epitope (p=0.029, 2-sided Fisher exact test).

### Maternal HIV variant neutralization sensitivity in paired plasma and breast milk

All transmission events studied in this cohort occurred postpartum. As breast milk is known to have lower neutralizing activity than plasma (37) and to be populated by locally-replicating variants (42), thus we sought to determine if bnAb resistance profiles were distinct between breast milk and plasma HIV variants in women with plasma bnAb activity. Autologous viruses from four CHAVI009 mothers, one transmitting and three non-transmitting, with plasma broad neutralizing activity were previously sequenced via single genome amplification (SGA) from plasma and breast milk (37). In transmitting mother 1209, *env* 1209bmH5 was identified as most similar to the infant 1209 virus population through phylogenetic analysis (Fig. 4) We tested these autologous Env pseudoviruses for bnAb sensitivity against monoclonal antibodies PGT121, PG9, VRC01, and 10E8 (Fig 5A). No notable differences in bnAb sensitivity of the milk and plasma viruses were found in the mothers tested. Pseudoviruses prepared using SGAs from both breast milk and plasma of mothers 1209 and 4707 were uniformly resistant to VRC01. Autologous viruses from mother 3305 were uniformly resistant to neutralization against bnAbs PGT121 and PG9. In summary, the neutralization sensitivity profiles of breast milk and plasma-isolated Envs appear to be comparable in this cohort.

**Figure 4.**
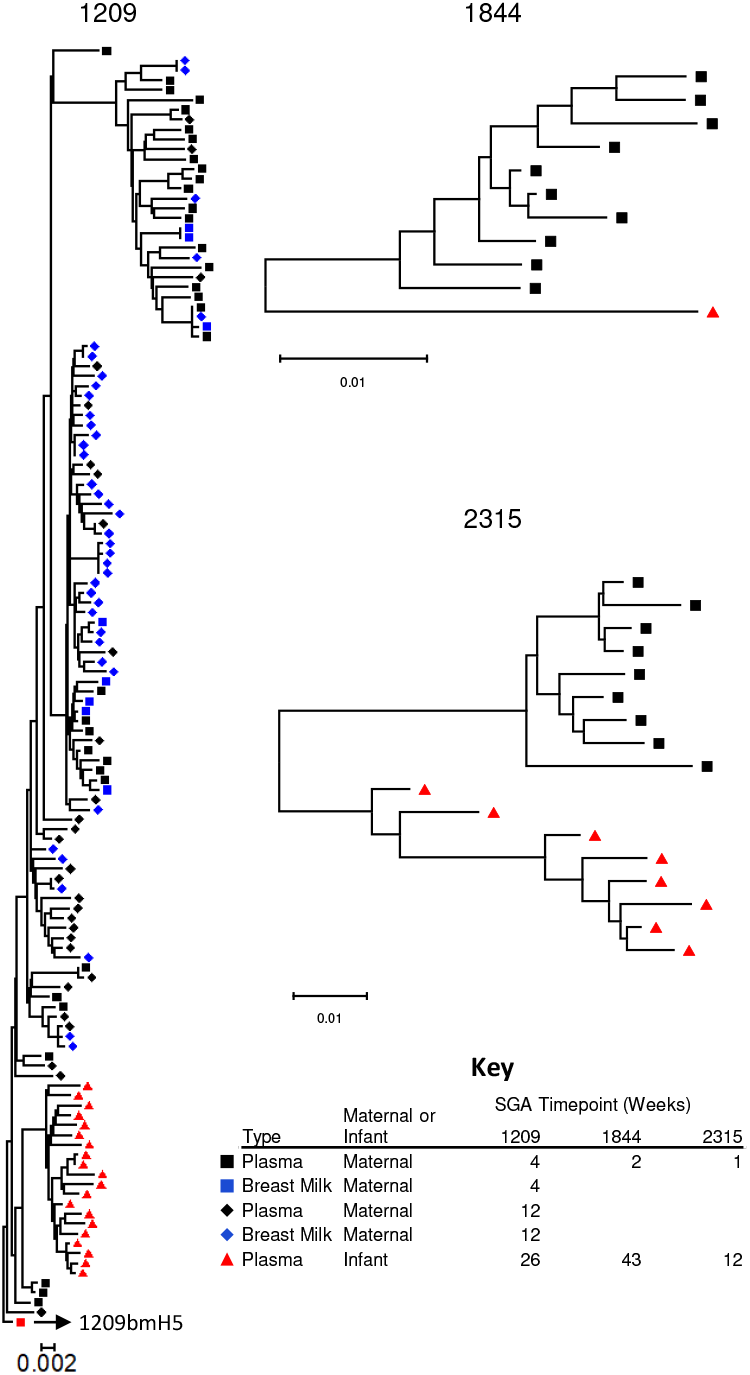
Phylogenetic tree analysis of env SGA from transmitting mothers and their infants. Phylogenetic trees were prepared using the Kimura parameter 2 method. The 1209 tree is rooted at maternal milk sequence 1209bmH5, the env most closely related to the infant virus population. Maternal SGA timepoints are indicated as weeks postpartum while infant SGA timepoints are indicated as weeks post infection.

**Figure 5.**
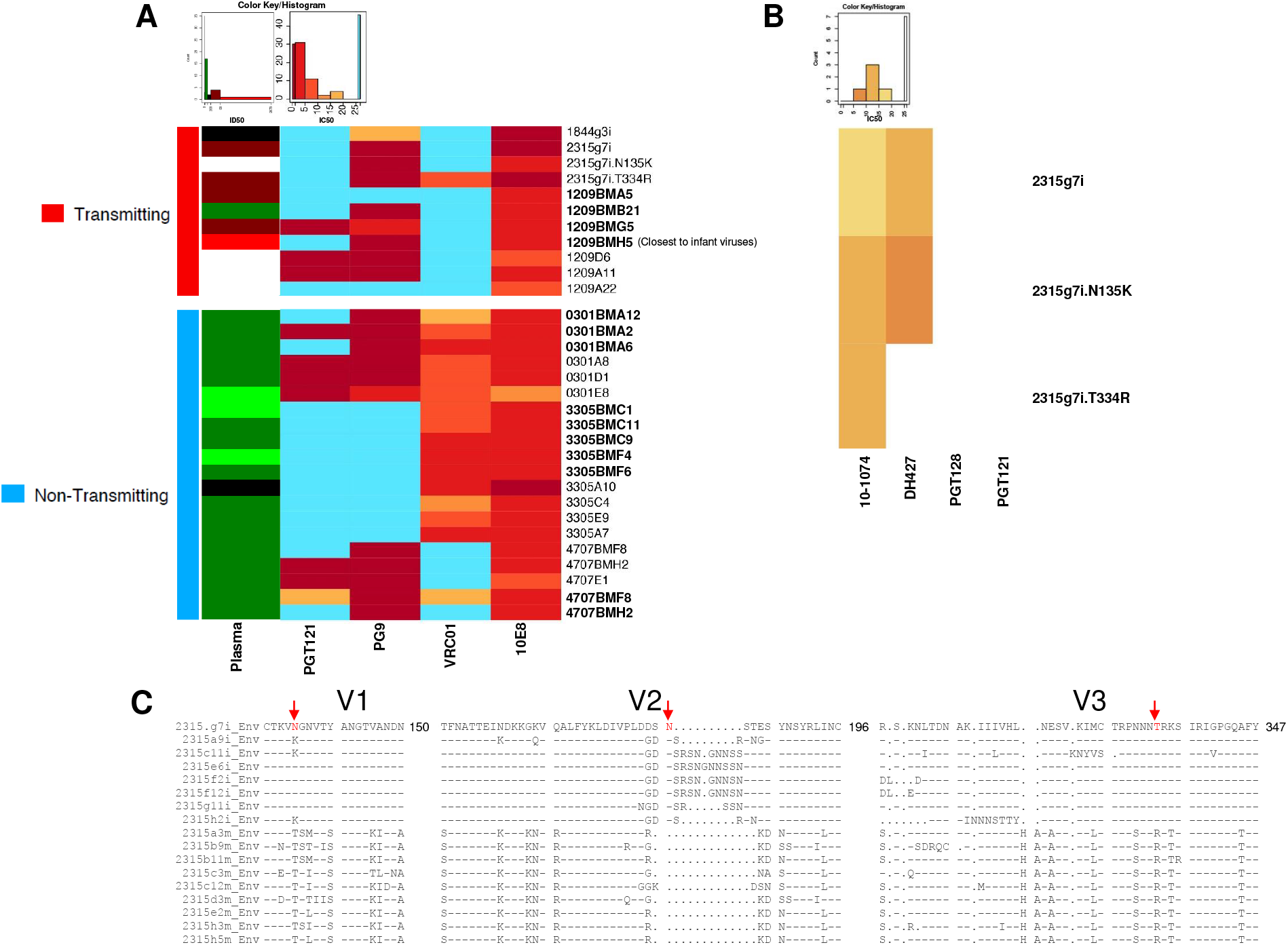
BnAb neutralization sensitivity of maternal and infant HIV Env pseudoviruses isolated from breast milk and plasma. (A) Neutralization sensitivity to autologous maternal plasma, PGT121 V3 glycan-specific IgG bnAb, PG9 V2 glycan-specific IgG bnAb, VRC01 CD4bs-specific IgG bnAb, and 10E8 MPER-specific IgG bnAb. Neutralization was measured using ID_50_ for plasma and IC_50_ for bnAbs as described in the color keys/histograms. Blood plasma isolated variants are denoted in normal font and Breast milk isolated variants are denoted with “BM” in the name and are in bold. (B) Neutralization sensitivity of 2315g7i autologous pseudovirus and mutants to V3 glycan-specific IgG bnAbs 10-1074, DH427, PGT128, and PGT121. (C) Transmitting mother-infant pair 2315 isolated HIV env sequences had mutations in the V1, V2, and V3 glycan regions that could affect V3 glycan-specific binding. Infant sequences end in “i” and maternal sequences end in “m”. The mutations observed were K135N or T, V2 loop extensions at 188, and T334R mutations, respectively as denoted with red arrows and text.

### VRC01-blocking activity of maternal plasma

To explore whether the bnAb resistance of circulating maternal HIV variants could be explained by bnAb activity that was unmappable in the mutant HIV Env pseudovirus assay, we measured VRC01-blocking activity in maternal plasma (Fig. 6). All HIV Env pseudoviruses from mothers 1209 and 4707 had uniform resistance to VRC01 (Fig. 5). To determine if VRC01-like antibody binding was present in plasma, we measured VRC01-blocking activity in maternal plasma using a competition ELISA, where percent blocking over 50 percent was considered positive (Fig. 6). Interestingly, some mothers (129, 1306, 0205, 3507, 2709, 4707) with unmappable plasma bnAb activity by neutralization assay had detectable levels of VRC01 blocking, while plasma of 1209, which was mapped to V2 glycan-specific neutralization activity and who postnatally-transmitted a VRC01-resistant virus, too had positive VRC01 blocking (68.6% blocking). As expected, autologous HIV Env pseudoviruses from mother 3305 were all sensitive to VRC01 and little VRC01 blocking activity was observed in plasma (14.1% blocking).

### Infant T/F variant sensitivity to maternal plasma neutralizing activity and mapped bnAb specificities

We sought to determine whether viruses transmitted to infants were escape variants of maternal plasma bnAb activity. We analyzed 3 mothers who transmitted virus to their infant, had plasma activity mapped to a defined epitope, and had successful identification of infant virus *env* sequences (1844i, 2315i, and 1209i). For infant 1209i, the viral population was too diverse to identify an infant T/F virus, so the phylogenetically closest maternal breast milk virus (1209bmH5) was used to represent the infant virus population(37). Phylogenetic tree analysis revealed varying levels of genetic similarity between maternal and infant virus envelopes (Fig. 4). A T/F could not be identified for 1844 and 2315 because of too few infant Envs and extensive diversity among the isolated Envs respectively. Since 1844i and 2315i Envs were genetically different from their maternal counterparts, we chose to focus on these infant sequences for neutralization sensitivity analysis.

We addressed whether infant viruses acquired mutations that would lead to neutralization escape from paired maternal plasma bnAbs. While subject 1209’s maternal plasma bnAb activity was mapped to the V2 glycan specificity, most of the autologous viruses from this sample, including 1209bmH5, were sensitive to V2 glycan-targeting bnAb PG9 and paired maternal plasma (Fig. 5A), indicating that this transmission was not related to escape from the dominant maternal bnAb response. 1844g3i was sensitive to paired maternal plasma neutralization and MPER-targeting bnAb 10E8 (Fig. 5A) yet neutralization resistant to all other bnAbs tested, including CD4bs-targeting VRC01. Interestingly,1844 maternal plasma was mapped to the N276 CD4 binding site. No mutations in infant virus 1844g3i *env* were identified in the N276 region, yet it was phenotypically resistant to VRC01. Both maternal and infant viruses did have an isoleucine insertion at position 460 which could affect the glycosylation of the G458 glycan, which is necessary for the binding of VRC01 and other bnAbs to the CD4bs (50–52).

Infant HIV Env pseudovirus 2315g7i was potently neutralized by paired maternal plasma, PG9, and 10E8 but was neutralization resistant to VRC01 and PGT121 (Fig. 5A). Interestingly, maternal plasma corresponding to infant 2315 was mapped to the N332 V3 glycan specificity and 2315g7i was resistant to corresponding bnAb PGT121 (Fig. 3, Fig. 5A). We explored this V3 glycan-targeting bnAb resistance of 2315g7i by creating Env pseudovirus mutants 2315g7i.N135K and 2315g7i.T334R. We looked at differences in the V1 and V2 regions of 2315 maternal and infant virus *envs*, as mutations in these regions of the envelope can affect V3-specific bnAb activity (53, 54). These regions in the infant virus *envs* vary greatly from the maternal variants (Fig. 5C). Notably, 5 of 8 infant viruses had a K135N mutation which led to the insertion of a putative glycosylation site. Reverting this mutation in 2315g7i.N135K was hypothesized to have increased neutralization resistance to V2 glycan and possibly V3 glycan-targeting bnAbs, but the mutant pseudovirus had a similar neutralization profile to the wild-type virus (Fig. 6A). For the V3 glycan region, all infant *envs* included the N332 V3 glycan while all maternal *env* sequences contained mutation T334R which deglycosylates the N332 V3 glycan region (Fig. 5C). We hypothesized that inducing mutation T334R in pseudovirus 2315g7i would increase the Env’s neutralization resistance to a larger number of V3 glycan-targeting bnAbs. 2315g7i.T334R still retained neutralization resistance to PGT121 and PGT128 and acquired neutralization resistance to DH427. (Fig. 5B). However, this mutation also slightly increased neutralization sensitivity to 10-1074 compared to the wild-type pseudovirus (1.96 fold).

**Figure 6.**
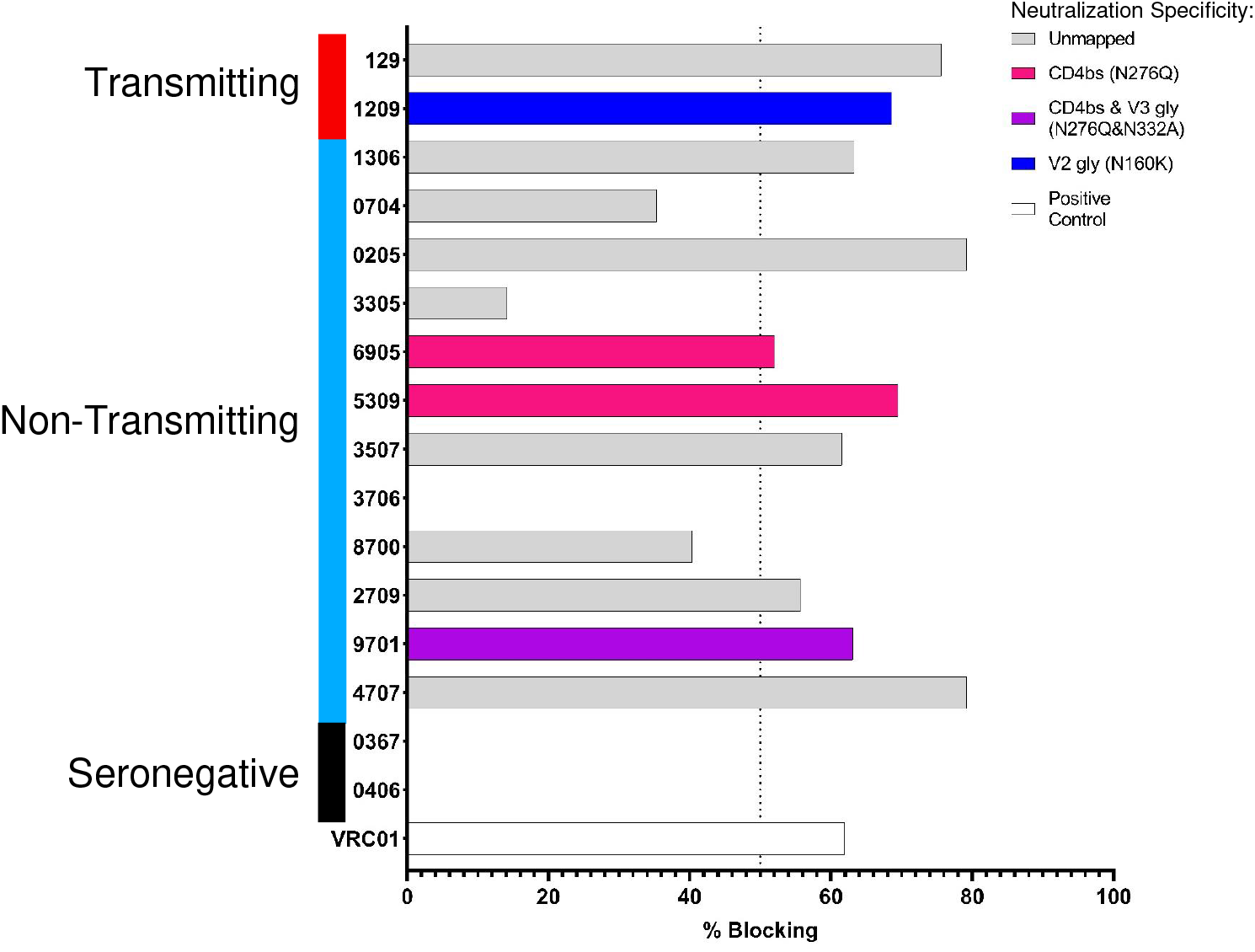
VRC01 bnAb-blocking activity of plasma of mothers with broad neutralizing activity. Percent VRC01 blocking was found by calculating the percent change in OD binding compared to uninhibited VRC01 binding to 1086Cgp120K160N. The dotted lines indicate the positivity cutoff of 50% blocking.

Finally, we compared the length and N-linked glycosylation in the V1V2 and V5 regions as well as the total glycosylation in maternal and corresponding infant envelopes (Fig. 7), as lengthening and glycosylation in these regions can affect bnAb neutralization sensitivity (53–55). Interestingly, infant 2315i *env* sequences had significantly higher glycosylation sites in the hypervariable V1V2 regions compared to maternal *env* sequences (p=0.00032 by 2-sided Wilcoxon test) and significantly longer V1V2 hypervariable loops compared to maternal sequences (p=0.0003 by 2-sided Wilcoxon test). Compared to maternal Envs, all 8 infant sequences have insertions 3-11 amino acid long at HXB2 position 188 (Fig. 5C). While previous studies have found that T/F viruses tend to have shorter V1V2 lengths as it favors transmission fitness(56), longer V1 V2 loops have been associated with resistance to V3 and CD4bs bnAbs (54, 55), consistent with the fact that these viruses are all neutralization resistant to PGT121 and VRC01 (Fig 5A). This suggests that different transmission fitness pressures could be working on infant T/F viruses in the context of MTCT vertical transmissions. Finally, the net charge of 2315i V1V2 regions were also significantly different than maternal variants. 2315i V1V2 regions tended to be negatively charged while maternal variants were predominantly positively charged (p=0.00044). It has been previously shown that a net positive charge in the V2 strand is correlated with sensitivity to PG9 and PG16 (57), however 2315g7i is phenotypically sensitive to PG9 (Fig. 5A). In summary, our study is suggestive, yet not definitive on whether infant *env* mutations are frequently driven by maternal bnAb selection pressure.

**Figure 7.**
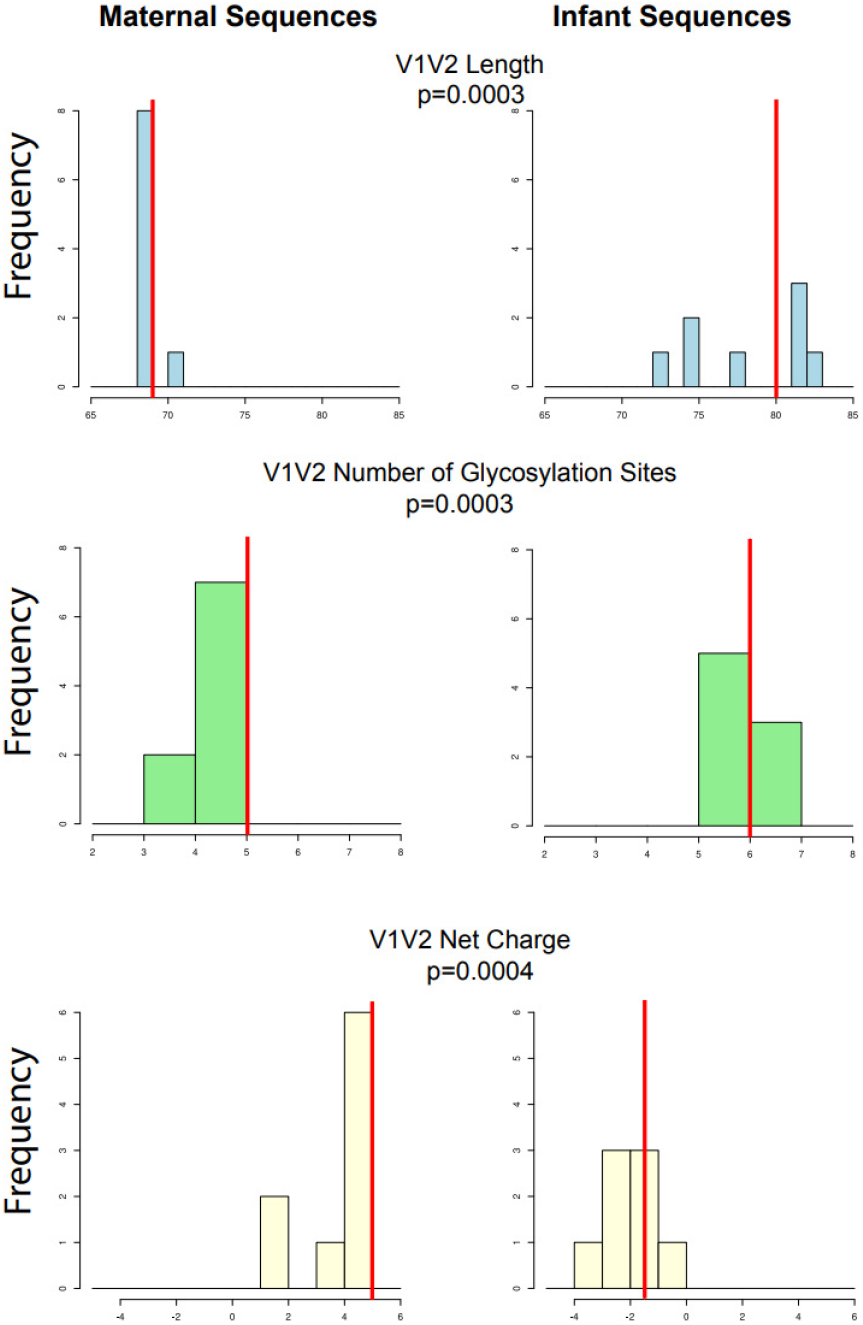
V1V2 variable characteristics comparing pair 2351 maternal and infant Envs. Frequency plots of the V1V2 variable region lengths (top), number of glycosylation sites (middle), and net charge (bottom) for 2351 maternal sequences (left) compared to 2351 infant sequences. All p-values were obtained using a 2-sided Wilcoxon test. Red bars indicate the medians of each distribution.

## Discussion

To achieve an HIV-free generation, supplemental strategies in conjunction with ART will be required to prevent MTCT. In this study, we investigated vertical HIV transmission in pregnant women with plasma neutralizing breadth and potency to understand the characteristics of epitope-specific responses that may be protective against virus transmission and/or contribute to selection of transmitted variants during postpartum transmission. Importantly, due to the limited success of the single bnAb prevention approach studied in the Antibody Mediated Prophylaxis (AMP) trial in adults (58), it is clear that understanding transmitted virus escape of bnAb activity is critical to advancing bnAb-based prophylaxis and therapy. The gap in knowledge of protective bnAb combinations for mothers and infants urgently needs addressing, as it will define vaccine strategies and antibody therapies that will be effective in preventing MTCT.

We aimed to define protective bnAb activity in postpartum transmitting and non-transmitting mothers from the BAN and CHAVI cohorts by mapping their epitope-specificity using a panel of mutant Env pseudoviruses. We found that more transmitting women had bnAb responses that mapped to 1 or 2 known bnAb specificities compared to non-transmitting women, where the majority were unable to be mapped to common bnAb epitopes, indicating multiple specificities of the neutralization response, responses against undefined epitopes, or responses that were too low to be detected in the pseudovirus neutralization epitope mapping assay. This pattern suggests that multiple bnAb specificities are more common in non-transmitting than transmitting women with broad neutralizing activity and may be required to prevent MTCT. Alternatively, this data could also suggest that a currently undefined and infrequent specificity of bnAbs may play a role in prevention of MTCT.

We previously demonstrated that autologous HIV viruses isolated from transmitting mothers and their infected infants were enriched with genetic motifs that were associated with resistance to bnAbs (59), suggesting that acquired plasma neutralization-resistance is a common feature of vertically transmitted variants. We also demonstrated that an in utero-transmitted virus from a mother with plasma V3 glycan-specific bnAb activity was selected for its ability to escape the mother’s dominant bnAb specificity (41). In this study we described a mother-infant transmission pair where the infant envs exhibited V1V2 variable loop characteristics associated with V3 and/or CD4bs bnAb resistance: higher number of glycosylation sites compared to maternal envs, longer V1V2 insertions, and negative net charge. Thus, it is possible that mothers with a single dominant plasma bnAb specificity could drive the selection of resistant viral variants that are in turn, more likely to be transmitted to their infants. An *in vitro* study using HIV strain BG505 showed that plasma neutralization breadth is moderately correlated to ease of escape via a single mutation (60). Our work demonstrating the potential for lower vertical transmission risk in women with broad, unmappable neutralization activity suggests that transmission may be averted by maternal bnAbs that target uncommon epitopes with a lower capacity for antibody escape via single mutations or with polyclonal bnAb activity. Viruses circulating in women with polyclonal and broad neutralization activity may be less likely to escape bnAbs compared to maternal plasma with a single, dominant specificity, impeding transmission by preventing escape variants.

We also tested breast milk and plasma autologous viruses from four mothers for neutralization resistance against bnAbs targeting common bnAb epitopes. We observed no differences in neutralization sensitivities between breast milk and plasma viruses within each mother. This is consistent with a study of 13 subjects from Zambia, which did not find a restriction in viral genotype in breast milk (61) and a previous CHAVI009 study which observed limited compartmentalization of breast milk viruses (62). In this study, several mothers had plasma virus populations with uniform neutralization resistance to some bnAbs. Mothers 1209 and 4707 had viruses that were all neutralization resistant to CD4bs-specfic bnAb VRC01. Interestingly, mothers with VRC01-resistant plasma viruses had detectable plasma VRC01-blocking activity not identified in the plasma bnAb mapping assay. This could be because binding assays are generally more sensitive than neutralization assays, but as a result we could not confirm these antibodies’ function or contribution to the selection of VRC01-resistant viruses.

We were also able to generate infant HIV Env pseudoviruses for transmitting mother-infant pairs with epitope-mapped bnAb responses (1209, 2315, 1844). We hypothesized that infant HIV Env variants would be resistant to bnAbs targeting the same epitopes as found in the paired maternal plasma. In fact, infant HIV Env variants generated from mother-infant pairs 2315 and 1844 were resistant to the maternal mapped specificities, V3 glycan and CD4bs respectively. Despite this, the paired maternal plasma was able to neutralize both infant viruses potently, likely due to other neutralizing antibodies targeting unique specificities present in the plasma. These results suggest that bnAb selection pressure could be driving the transmission of bnAb-resistant viruses, but neutralization resistance may not be required for MTCT.

Our study is not without limitations. Our cohort is small due to the sample availability and neutralization activity selection criteria, as bnAb activity is detected in a minority of HIV-infected adults. Additionally, we isolated only a few maternal and infant *env* variants from the transmitting mother-infant pairs from timepoints late after transmission due to lack of sample availability. Thus, we were not always able to determine if infant-transmitted variants were resistant to paired maternal antibody specificities, as the sequenced virus populations could experience reversions when there is less maternal antibody selection pressure. Another limitation of this study is that we only focused on postpartum transmissions, so observations in this study may not be extrapolated to other modes of MTCT. Future studies should expand our understanding of protective bnAbs by investigating all modes of MTCT, increasing sample size, increasing autologous viruses isolated, and characterizing maternal monoclonal antibodies. Despite our study limitations, this is to our knowledge the largest study identifying differences in bnAb specificity of maternal plasma between transmitting and non-transmitting women.

Overall, our results demonstrate that transmitting women with plasma broad neutralizing activity often have antibody activity directed against dominant bnAb epitopes, compared to non-transmitting women whose broad neutralizing activity was unable to be mapped to known specificities. Thus, possessing only a single, common bnAb specificity may be a risk factor for vertical HIV transmission. These results have important implications regarding the use of passive bnAb administration or bnAb-targeting vaccines in HIV-infected pregnant women, who may require the administration or induction of two or more bnAb specificities to prevent maternal virus variants from acquiring antibody escape mutations that would allow for vertical transmission in the setting of pre-existing antibodies. This work underscores supplemental strategies that administer bnAbs to mothers and infants to prevent MTCT must be developed as combinations to effectively contain vertical transmission of HIV and make further gains in the elimination of pediatric HIV.

## Acknowledgements

We are grateful to the women and their infants who participated in the BAN and CHAVI009 studies and provided us with an invaluable resource. We also thank Kshitij Wagh for helpful discussions. Manish Sagar is supported by NIH grant AI122209 (M.S.), K24-AI145661, and P30 - AI042853. Sallie R. Permar is supported by NIH NIAID grants DP2 HD075699, R01 AI106380, and P01 AI117915. This work was also supported by NIAID grant R01 AI122909, awarded to Feng Gao and Sallie R. Permar. The funders had no role in study design, data collection, decision for publication, or preparation of the manuscript. The content is solely the view of the authors and does not necessarily represent the official views of the National Institutes of Health or Centers for Disease Control and Prevention.

## Works Cited

1. UNAIDS. Global HIV & AIDS statistics — 2021 Fact sheet [Available from: https://www.unaids.org/en/resources/fact-sheet.

2. Linnemayr S, Jennings Mayo-Wilson L, Saya U, Wagner Z, MacCarthy S, Walukaga S, et al. HIV Care Experiences During the COVID-19 Pandemic: Mixed-Methods Telephone Interviews with Clinic-Enrolled HIV-Infected Adults in Uganda. AIDS Behav. 2021;25(1):28–39.

3. John GC, Kreiss J. Mother-to-child transmission of human immunodeficiency virus type 1. Epidemiol Rev. 1996;18(2):149–57.

4. Teasdale CA, Marais BJ, Abrams EJ. HIV: prevention of mother-to-child transmission. BMJ Clin Evid. 2011;2011.

5. Wolinsky SM, Wike CM, Korber BT, Hutto C, Parks WP, Rosenblum LL, et al. Selective transmission of human immunodeficiency virus type-1 variants from mothers to infants. Science. 1992;255(5048):1134–7.

6. Ahmad N, Baroudy BM, Baker RC, Chappey C. Genetic-Analysis of Human-Immunodeficiency-Virus Type-1 Envelope V3 Region Isolates from Mothers and Infants after Perinatal Transmission. Journal of Virology. 1995;69(2):1001–12.

7. Dickover RE, Garratty EM, Plaeger S, Bryson YJ. Perinatal transmission of major, minor, and multiple maternal human immunodeficiency virus type 1 variants in utero and intrapartum. J Virol. 2001;75(5):2194–203.

8. Scarlatti G, Leitner T, Hodara V, Halapi E, Rossi P, Albert J, et al. Neutralizing antibodies and viral characteristics in mother-to-child transmission of HIV-1. AIDS. 1993;7:S45–8.

9. Zhang H, Orti G, Du Q, He J, Kankasa C, Bhat G, et al. Phylogenetic and phenotypic analysis of HIV type 1 env gp120 in cases of subtype C mother-to-child transmission. AIDS Res Hum Retroviruses. 2002;18(18):1415–23.

10. Zhang H, Tully DC, Hoffmann FG, He J, Kankasa C, Wood C. Restricted genetic diversity of HIV-1 subtype C envelope glycoprotein from perinatally infected Zambian infants. PLoS One. 2010;5(2):e9294.

11. Garber DA, Guenthner P, Mitchell J, Ellis S, Gazumyan A, Nason M, et al. Broadly neutralizing antibody-mediated protection of macaques against repeated intravenous exposures to simian-human immunodeficiency virus. AIDS. 2021;35(10):1567–74.

12. Saunders KO, Pegu A, Georgiev IS, Zeng M, Joyce MG, Yang ZY, et al. Sustained Delivery of a Broadly Neutralizing Antibody in Nonhuman Primates Confers Long-Term Protection against Simian/Human Immunodeficiency Virus Infection. J Virol. 2015;89(11):5895–903.

13. Saunders KO, Wang L, Joyce MG, Yang ZY, Balazs AB, Cheng C, et al. Broadly Neutralizing Human Immunodeficiency Virus Type 1 Antibody Gene Transfer Protects Nonhuman Primates from Mucosal Simian-Human Immunodeficiency Virus Infection. J Virol. 2015;89(16):8334–45.

14. Rosenberg YJ, Jiang X, Cheever T, Coulter FJ, Pandey S, Sack M, et al. Protection of Newborn Macaques by Plant-Derived HIV Broadly Neutralizing Antibodies: a Model for Passive Immunotherapy during Breastfeeding. J Virol. 2021;95(18):e0026821.

15. Julg B, Tartaglia LJ, Keele BF, Wagh K, Pegu A, Sok D, et al. Broadly neutralizing antibodies targeting the HIV-1 envelope V2 apex confer protection against a clade C SHIV challenge. Sci Transl Med. 2017;9(406).

16. Xu L, Pegu A, Rao E, Doria-Rose N, Beninga J, McKee K, et al. Trispecific broadly neutralizing HIV antibodies mediate potent SHIV protection in macaques. Science. 2017;358(6359):85–90.

17. Kumar A, Smith CEP, Giorgi EE, Eudailey J, Martinez DR, Yusim K, et al. Infant transmitted/founder HIV-1 viruses from peripartum transmission are neutralization resistant to paired maternal plasma. PLoS Pathog. 2018;14(4):e1006944.

18. Russell ES, Kwiek JJ, Keys J, Barton K, Mwapasa V, Montefiori DC, et al. The genetic bottleneck in vertical transmission of subtype C HIV-1 is not driven by selection of especially neutralization-resistant virus from the maternal viral population. J Virol. 2011;85(16):8253–62.

19. Wu X, Parast AB, Richardson BA, Nduati R, John-Stewart G, Mbori-Ngacha D, et al. Neutralization escape variants of human immunodeficiency virus type 1 are transmitted from mother to infant. J Virol. 2006;80(2):835–44.

20. Dickover R, Garratty E, Yusim K, Miller C, Korber B, Bryson Y. Role of maternal autologous neutralizing antibody in selective perinatal transmission of human immunodeficiency virus type 1 escape variants. J Virol. 2006;80(13):6525–33.

21. Milligan C, Omenda MM, Chohan V, Odem-Davis K, Richardson BA, Nduati R, et al. Maternal Neutralization-Resistant Virus Variants Do Not Predict Infant HIV Infection Risk. mBio. 2016;7(1):e02221–15.

22. Lynch JB, Nduati R, Blish CA, Richardson BA, Mabuka JM, Jalalian-Lechak Z, et al. The breadth and potency of passively acquired human immunodeficiency virus type 1-specific neutralizing antibodies do not correlate with the risk of infant infection. J Virol. 2011;85(11):5252–61.

23. deCamp A, Hraber P, Bailer RT, Seaman MS, Ochsenbauer C, Kappes J, et al. Global panel of HIV-1 Env reference strains for standardized assessments of vaccine-elicited neutralizing antibodies. J Virol. 2014;88(5):2489–507.

24. Ghulam-Smith M, Olson A, White LF, Chasela CS, Ellington SR, Kourtis AP, et al. Maternal but Not Infant Anti-HIV-1 Neutralizing Antibody Response Associates with Enhanced Transmission and Infant Morbidity. mBio. 2017;8(5).

25. Stamatatos L, Morris L, Burton DR, Mascola JR. Neutralizing antibodies generated during natural HIV-1 infection: good news for an HIV-1 vaccine? Nat Med. 2009;15(8):866–70.

26. Binley JM, Lybarger EA, Crooks ET, Seaman MS, Gray E, Davis KL, et al. Profiling the specificity of neutralizing antibodies in a large panel of plasmas from patients chronically infected with human immunodeficiency virus type 1 subtypes B and C. J Virol. 2008;82(23):11651–68.

27. Doria-Rose NA, Klein RM, Manion MM, O’Dell S, Phogat A, Chakrabarti B, et al. Frequency and phenotype of human immunodeficiency virus envelope-specific B cells from patients with broadly cross-neutralizing antibodies. J Virol. 2009;83(1):188–99.

28. Li Y, Svehla K, Louder MK, Wycuff D, Phogat S, Tang M, et al. Analysis of neutralization specificities in polyclonal sera derived from human immunodeficiency virus type 1-infected individuals. J Virol. 2009;83(2):1045–59.

29. Sather DN, Armann J, Ching LK, Mavrantoni A, Sellhorn G, Caldwell Z, et al. Factors associated with the development of cross-reactive neutralizing antibodies during human immunodeficiency virus type 1 infection. J Virol. 2009;83(2):757–69.

30. Simek MD, Rida W, Priddy FH, Pung P, Carrow E, Laufer DS, et al. Human immunodeficiency virus type 1 elite neutralizers: individuals with broad and potent neutralizing activity identified by using a high-throughput neutralization assay together with an analytical selection algorithm. J Virol. 2009;83(14):7337–48.

31. Hraber P, Seaman MS, Bailer RT, Mascola JR, Montefiori DC, Korber BT. Prevalence of broadly neutralizing antibody responses during chronic HIV-1 infection. Aids. 2014;28(2):163–9.

32. Wei X, Decker JM, Wang S, Hui H, Kappes JC, Wu X, et al. Antibody neutralization and escape by HIV-1. Nature. 2003;422(6929):307–12.

33. Bosch KA, Rainwater S, Jaoko W, Overbaugh J. Temporal analysis of HIV envelope sequence evolution and antibody escape in a subtype A-infected individual with a broad neutralizing antibody response. Virology. 2010;398(1):115–24.

34. Dingens AS, Haddox HK, Overbaugh J, Bloom JD. Comprehensive Mapping of HIV-1 Escape from a Broadly Neutralizing Antibody. Cell Host Microbe. 2017;21(6):777–87 e4.

35. Liao HX, Lynch R, Zhou T, Gao F, Alam SM, Boyd SD, et al. Co-evolution of a broadly neutralizing HIV-1 antibody and founder virus. Nature. 2013;496(7446):469–76.

36. Moore PL, Gray ES, Wibmer CK, Bhiman JN, Nonyane M, Sheward DJ, et al. Evolution of an HIV glycan-dependent broadly neutralizing antibody epitope through immune escape. Nat Med. 2012;18(11):1688–92.

37. Fouda GG, Mahlokozera T, Salazar-Gonzalez JF, Salazar MG, Learn G, Kumar SB, et al. Postnatally-transmitted HIV-1 Envelope variants have similar neutralization-sensitivity and function to that of nontransmitted breast milk variants. Retrovirology. 2013;10:3.

38. Nakamura KJ, Heath L, Sobrera ER, Wilkinson TA, Semrau K, Kankasa C, et al. Breast milk and in utero transmission of HIV-1 select for envelope variants with unique molecular signatures. Retrovirology. 2017;14(1):6.

39. Russell ES, Ojeda S, Fouda GG, Meshnick SR, Montefiori D, Permar SR, et al. Short communication: HIV type 1 subtype C variants transmitted through the bottleneck of breastfeeding are sensitive to new generation broadly neutralizing antibodies directed against quaternary and CD4-binding site epitopes. AIDS Res Hum Retroviruses. 2013;29(3):511–5.

40. Mabuka J, Goo L, Omenda MM, Nduati R, Overbaugh J. HIV-1 maternal and infant variants show similar sensitivity to broadly neutralizing antibodies, but sensitivity varies by subtype. AIDS. 2013;27(10):1535–44.

41. Martinez DR, Tu JJ, Kumar A, Mangold JF, Mangan RJ, Goswami R, et al. Maternal Broadly Neutralizing Antibodies Can Select for Neutralization-Resistant, Infant-Transmitted/Founder HIV Variants. mBio. 2020;11(2).

42. Salazar-Gonzalez JF, Salazar MG, Learn GH, Fouda GG, Kang HH, Mahlokozera T, et al. Origin and Evolution of HIV-1 in Breast Milk Determined by Single-Genome Amplification and Sequencing. Journal of Virology. 2011;85(6):2751–63.

43. Sacha CR, Vandergrift N, Jeffries TL, McGuire E, Fouda GG, Liebl B, et al. Restricted isotype, distinct variable gene usage, and high rate of gp120 specificity of HIV-1 envelope-specific B cells in colostrum compared with those in blood of HIV-1-infected, lactating African women. Mucosal Immunology. 2015;8(2):316–26.

44. Chasela CS, Hudgens MG, Jamieson DJ, Kayira D, Hosseinipour MC, Kourtis AP, et al. Maternal or Infant Antiretroviral Drugs to Reduce HIV-1 Transmission. New England Journal of Medicine. 2010;362(24):2271–81.

45. Wagh K, Bhattacharya T, Williamson C, Robles A, Bayne M, Garrity J, et al. Optimal Combinations of Broadly Neutralizing Antibodies for Prevention and Treatment of HIV-1 Clade C Infection. PLoS Pathog. 2016;12(3):e1005520.

46. Sarzotti-Kelsoe M, Bailer RT, Turk E, Lin C-l, Bilska M, Greene KM, et al. Optimization and validation of the TZM-bl assay for standardized assessments of neutralizing antibodies against HIV-1. J Immunol Methods. 2014;409:131–46.

47. Saunders KO, Lee E, Parks R, Martinez DR, Li D, Chen H, et al. Neutralizing antibody vaccine for pandemic and pre-emergent coronaviruses. Nature. 2021;594(7864):553–9.

48. Gouy M, Guindon S, Gascuel O. SeaView Version 4: A Multiplatform Graphical User Interface for Sequence Alignment and Phylogenetic Tree Building. Molecular Biology and Evolution. 2010;27(2):221–4.

49. Kumar S, Stecher G, Li M, Knyaz C, Tamura K. MEGA X: Molecular Evolutionary Genetics Analysis across Computing Platforms. Mol Biol Evol. 2018;35(6):1547–9.

50. Chuang G-Y, Liou D, Kwong PD, Georgiev IS. NEP: web server for epitope prediction based on antibody neutralization of viral strains with diverse sequences. Nucleic Acids Research. 2014;42(W1):W64–W71.

51. Wu X, Zhou T, Zhu J, Zhang B, Georgiev I, Wang C, et al. Focused Evolution of HIV-1 Neutralizing Antibodies Revealed by Structures and Deep Sequencing. Science. 2011;333(6049):1593–602.

52. Zhou T, Georgiev I, Wu X, Yang ZY, Dai K, Finzi A, et al. Structural Basis for Broad and Potent Neutralization of HIV-1 by Antibody VRC01. Science. 2010;329(5993):811–7.

53. Sagar M, Wu X, Lee S, Overbaugh J. Human Immunodeficiency Virus Type 1 V1-V2 Envelope Loop Sequences Expand and Add Glycosylation Sites over the Course of Infection, and These Modifications Affect Antibody Neutralization Sensitivity. Journal of Virology. 2006;80(19):9586–98.

54. Registre L, Moreau Y, Ataca ST, Pulukuri S, Henrich TJ, Lin N, et al. HIV-1 Coreceptor Usage and Variable Loop Contact Impact V3 Loop Broadly Neutralizing Antibody Susceptibility. J Virol. 2020;94(2).

55. Bricault CA, Yusim K, Seaman MS, Yoon H, Theiler J, Giorgi EE, et al. HIV-1 Neutralizing Antibody Signatures and Application to Epitope-Targeted Vaccine Design. Cell Host Microbe. 2019;25(1):59–72.e8.

56. Kariuki SM, Selhorst P, Ariën KK, Dorfman JR. The HIV-1 transmission bottleneck. Retrovirology. 2017;14(1):22.

57. Ringe R, Phogat S, Bhattacharya J. Subtle alteration of residues including N-linked glycans in V2 loop modulate HIV-1 neutralization by PG9 and PG16 monoclonal antibodies. Virology. 2012;426(1):34–41.

58. Data from Antibody-Mediated Prevention Studies Advance the Field and Show the Challenges that Lie Ahead [press release]. 01/26/2021 2021.

59. Kumar A, Giorgi EE, Tu JJ, Martinez DR, Eudailey J, Mengual M, et al. Mutations that confer resistance to broadly-neutralizing antibodies define HIV-1 variants of transmitting mothers from that of non-transmitting mothers. medRxiv. 2021:2021.01.07.21249396.

60. Dingens AS, Arenz D, Weight H, Overbaugh J, Bloom JD. An Antigenic Atlas of HIV-1 Escape from Broadly Neutralizing Antibodies Distinguishes Functional and Structural Epitopes. Immunity. 2019;50(2):520–32 e3.

61. Heath L, Conway S, Jones L, Semrau K, Nakamura K, Walter J, et al. Restriction of HIV-1 Genotypes in Breast Milk Does Not Account for the Population Transmission Genetic Bottleneck That Occurs following Transmission. PLOS ONE. 2010;5(4):e10213.

62. Salazar-Gonzalez JF, Salazar MG, Learn GH, Fouda GG, Kang HH, Mahlokozera T, et al. Origin and evolution of HIV-1 in breast milk determined by single-genome amplification and sequencing. J Virol. 2011;85(6):2751–63.

